# Ectopic head regeneration after nervous system ablation in a sea anemone

**DOI:** 10.1101/2025.02.04.636403

**Authors:** Fatemeh Mazloumi Gavgani, Johanna E.M. Kraus, Joshua November, Layla Al-Shaer, Anna Cosima Seybold, Benjamin Lerstad, Harald Hausen, Michael J. Layden, Fabian Rentzsch

## Abstract

Some animals are able to regenerate all missing cell types and large body parts after bisection, a phenomenon called whole-body regeneration. Many of these animals regenerate the correct tissues and structures with remarkable fidelity according to the original polarity of the body, reflecting positional information present in the remaining tissue. Understanding the cellular and molecular basis of this positional information is a central question in regeneration biology. In planarians and acoels, muscle cells have been shown to carry such positional information, but where this information originates and whether this function is conserved in other highly regenerative animals, is not well understood. Here we use the cnidarian *Nematostella vectensis* to address the role of the nervous system in whole-body regeneration. We generated a transgenic line for conditional ablation of neurons and first showed that *Nematostella* can repeatedly regenerate its nervous system. Bisection experiments following nervous system ablation showed that all head fragments regenerate a second head instead of a foot, whereas foot fragments correctly regenerate the missing head. We further found that regenerating head fragments of nervous system-ablated animals increase the expression of Wnt signaling genes that in wildtype animals are only upregulated in regenerating foot fragments. These molecular changes and the initiation of ectopic head regeneration precede the re-appearance of neurons, suggesting that the nervous system does not directly control whether a head or foot will be regenerated. Instead, we propose a model in which the nervous system provides positional information to the tissue of the body column, and that this information allows foot regeneration by suppressing a default program for head regeneration.

## INTRODUCTION

Regeneration describes the ability to replace missing parts of the body after unexpected loss due to traumatic, toxic or pathogenic insults. It occurs at different scales, ranging from the replacement of specific cell types to the regrowth of large structures following amputation. The ability to regenerate varies greatly among animals, with some species being able to rebuild fully functional bodies with correct proportions from small pieces of tissue. This remarkable phenomenon is referred to as whole-body regeneration, and it has been described in planarians, acoels, cnidarians and other taxa [1–6]. Whole-body regeneration occurs with high precision according to the original polarity of the tissue or body fragment, i.e. the tail fragment regenerates a new head, and the head fragment a new tail. The genes and cells contributing to the formation of new tissues during the regeneration process have been studied in some detail, in particular in planarians [7–12]. It is less clear, however, in which cells or tissues the positional information resides that instructs the regeneration of the correct body part, head or tail. In planarians and acoels, subepidermal muscle cells express position control genes that regulate axial polarity during regeneration [13–15], but it remains unknown whether other tissues or cell types are involved in regulating the position-specific expression of these genes. In *Hydra*, positional information is present in epithelial cells, as concurrent elimination of other cell types does not interfere with correct head and foot regeneration [16, 17].

Here we report an unexpected role for the nervous system during regeneration in the sea anemone *Nematostella vectensis*. *Nematostella* belongs to Cnidaria, a group of animals whose remarkable regeneration abilities have been studied extensively [18–21]. The body of cnidarian polyps resembles a tube with one opening at the oral end that is surrounded by tentacles for capturing prey. The tissue at this opening is traditionally called the head and the opposite end of the body axis is called the foot or the aboral region. As other cnidarian polyps, *Nematostella* readily regenerates missing body parts after transverse bisection [22–27]. To address the role of the nervous system in regeneration, we generated an *elav1::nitroreductase-cerulean* transgenic line to specifically ablate a large fraction of the epidermal and gastrodermal neurons. Following bisection of ablated animals, we observed head regeneration in 100% of both head and tail fragments, demonstrating a key role for the nervous system in controlling axial polarity during whole-body regeneration.

## RESULTS

### Nervous system regeneration following conditional cell ablation

To test a possible role of the nervous system in *Nematostella* regeneration, we generated double transgenic animals expressing a bacterial *nitroreductase* (*ntr*) gene [28] and a separate *mOrange* reporter transgene under the control of the neuron-specific *Nematostella elav1* promoter (*elav1::ntr-cerulean, elav1::mOrange*). Expression from the *elav1* promoter covers large parts of both the epidermal and gastrodermal *Nematostella* nervous system [29].

We ablated the *elav1^+^* neurons of six week-old juvenile polyps by 72 h exposure to Nifurpirinol (Nfp), which is processed by Nitroreductase into a cytotoxic metabolite [30], and monitored regeneration of the neurons by the *elav1::mOrange* reporter transgene. Confocal microscopy suggested a near-complete absence of *elav1::mOrange^+^* neurons at the end of the treatment period (day 0, Fig. 1A, B) and electron microscopy confirmed the loss of neurite bundles along the parietal muscles of the body wall (Fig. 1C-F). To test the specificity of the ablation, we crossed the *elav1::ntr-cerulean* animals to other reporter lines. Neither *foxQ2d::mOrange* cells (a non-overlapping population of sensory cells, [31]) nor *MyHC::mCherry* cells (retractor muscle, [32]) were affected by ablation of the *elav1*-expressing neurons (Fig 1G-J). During regeneration, the first neurons were detectable 4 days after washout of Nfp and after 25 days the proportion of *elav1::mOrange* neurons reached the pre-ablation level (Fig. 1K-R). We conclude that *Nematostella* is capable of regenerating the nervous system in a wound-free experimental paradigm.

**Figure 1:**
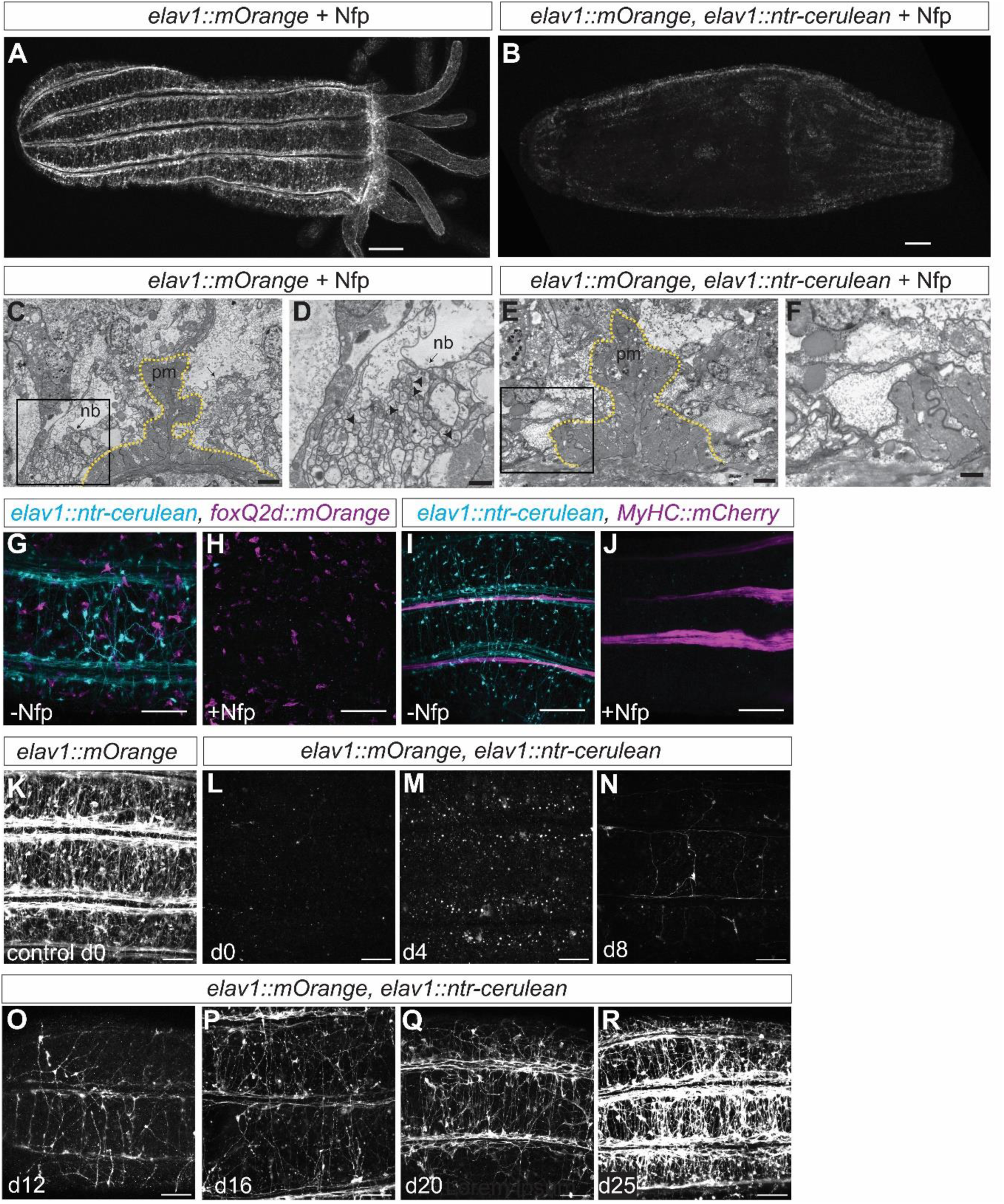
Regeneration of the nervous system in *Nematostella*. (A, B) Confocal microscopy images of six-week-old polyps after 72 h exposure to Nifurpirinol (Nfp), genotype is indicated at the top. The *elav1::mOrange* transgene labels a large part of the nervous system and is almost completely absent in animals containing the *elav1::ntr-cerulean* transgene (B). (C-F) Electron microscopy images of cross-sections of the body column after Nfp treatment in non-ablated (C, D) and ablated (E, F) animals. The parietal muscle (pm) at the base of one of the mesenteries is outlined by the yellow dashed line. (D, F) Higher magnification of the areas boxed in (C) and (E) shows that the longitudinal neurite bundles (nb) are absent in the ablated animal (E). Arrowheads in (D) indicate putative synaptic vesicles. (G-J) Confocal images of double transgenics show that ablation is restricted to the targeted cell population. Genotype on top, *elav1::ntr.cerulean* in turquoise, *foxQ2d::mOrange* (G, H) and *MyHC::mCherry* (I, J) in magenta. (K-R) Representative confocal images illustrating the time course of nervous system regeneration at approximately 50% body length along the oral-aboral axis. (K) is an animal treated with Nfp, but without the *elav1::ntr-cerulean* transgene. The first neurites are visible on day 4 (M). Scale bars: A, B, I, J 100µm; C, E 2µm; D, F 1µm; G, H, K-R 50µm

### Ablation of *elav1^+^* neurons reduces body wall motility but does not prevent feeding

Control animals (*elav1::mOrange*) treated with Nfp displayed a shorter body column and shorter tentacles (Fig. 2A, B). Following ablation of the *elav1^+^* neurons (*elav1::mOrange*, *elav1::ntr-cerulean* double transgenics), the body column of the animals appeared further shortened and they mostly kept the tentacles retracted into the gastric cavity (Fig. 2C). When presented with food (*Artemia* nauplii) they were able to extend the tentacles and capture prey at a low rate (Video S1). We speculate that this ability is based on neurons that are not ablated by the *elav1::ntr-cerulean* transgene. Unperturbed polyps display body wall constrictions that travel in oral to aboral direction (“peristaltic waves” [33], Video S2), but we did not observe such traveling constrictions in the first six days after ablation of the *elav1*^+^ neurons (Fig. 2D, E). Despite the reduced spontaneous motility, the ability to contract along the oral-aboral axis when touched at the oral pole was not impaired (Fig. 2F, Videos S3, 4).

**Figure 2.**
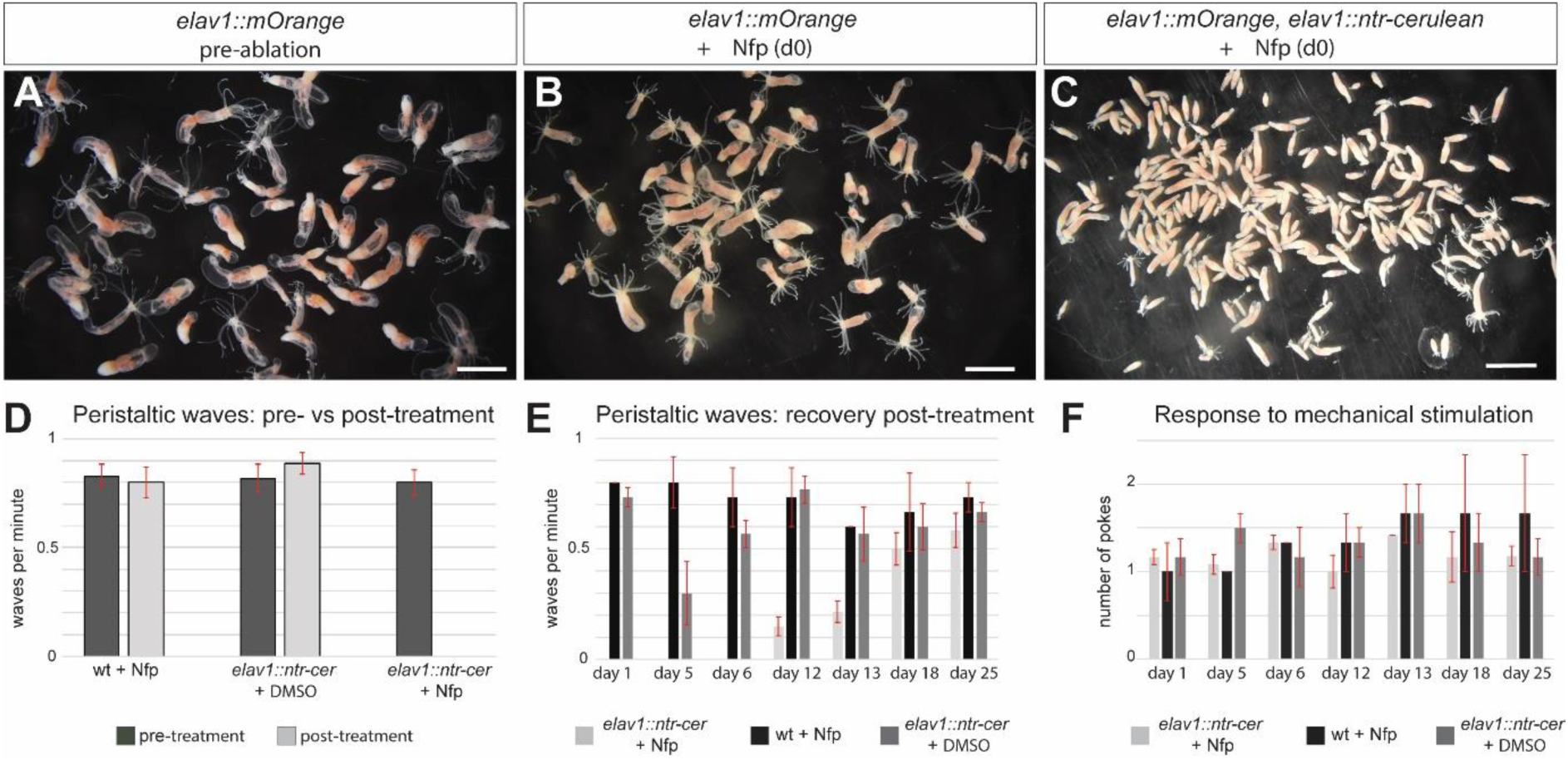
Impaired spontaneous activity after ablation of *elav1^+^*neurons. (A-C) Overview images of untreated polyps (A), Nfp-treated control polyps (B), and ablated Nfp-treated polyps carrying the *elav1::ntr-cer* transgene (C). After ablation, polyps keep the tentacles mainly retracted into the gastric cavity (C). (D) Quantification of the number of peristaltic waves per minute along the body column before (dark grey) and after (light grey) treatment. Animals were recorded for five minutes before and at day 1 following treatment, the number of waves per minute is shown. Genotype and treatment are indicated below the bars, n = 21-22 per treatment. Peristaltic waves are absent only after ablation of the *elav1^+^*neurons. (E) Quantification of the recovery of peristaltic waves during regeneration of the *elav1^+^* nervous system. Peristaltic waves re-appeared 12 days after ablation. n = 15 (*elav1::ntr-cer* + Nfp), 10 (*elav1::ntr-cer* + DMSO and wt + Nfp). (F) Quantification of contraction in response to poking during regeneration. The plot shows the average number of pokes required for eliciting contraction. No significant difference is observed in ablated vs non-ablated animals. n = 15 (*elav1::ntr-cer* + Nfp), 10 (*elav1::ntr-cer* + DMSO and wt + Nfp). Scale bars in (A-C) 3mm.

### Head fragments regenerate a second head after nervous system ablation

To address a possible role of the nervous system in whole body regeneration, we ablated the *elav1^+^* neurons (*elav1::mOrange*, *elav1::ntr-cerulean* double transgenics), washed out the Nifurpirinol and bisected the polyps at 50% body length. We observed that all foot fragments regenerated a new head with the first tentacles being visible 3 days post amputation (dpa), similar to control animals (*elav1::mOrange* treated with Nfp, data not shown). To our surprise, and in contrast to control animals, all head fragments of ablated animals regenerated a second head with tentacles within 6 dpa (Fig. 3A-D). The animals remained double-headed for up to several months and eventually separated into two polyps by transverse fission. The first ectopic heads became visible at 3 dpa, at a time when no or only very few regenerated *elav1::mOrange^+^* neurons can be observed. To test further whether the regeneration of ectopic heads correlates with the absence of the nervous system, we re-bisected the double-headed polyps at different times after the first bisection. We found that the number of animals regenerating a second head decreased with increasing time after the first bisection (Fig 3E). When re-bisected 6 days after the first bisection (and thus 6 days after washout of Nfp), 75 ± 5% of the fragments regenerated a second head (Fig. 3). This number dropped to 44.8 ± 4.7% at 13 days after the first bisection, and to 5 ± 5.6% at 25 days (Fig. 3E). Ectopic head regeneration thus preceded the regeneration of *elav1^+^* neurons and became less frequent when the number of these neurons gradually returned to wild type levels.

**Figure 3.**
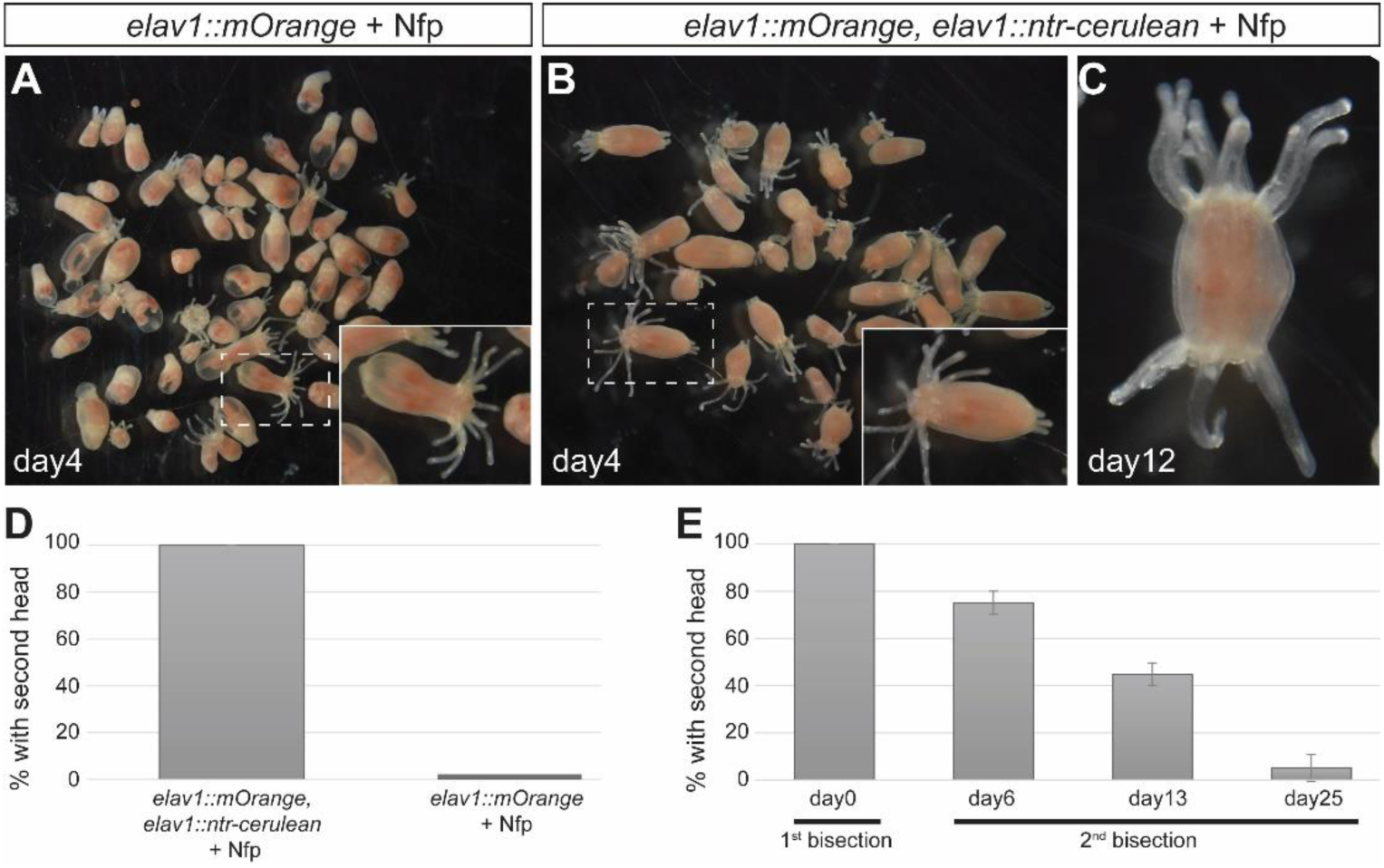
Ectopic head regeneration after ablation of the *elav1^+^* neurons. (A, B) Overview images of ablated (A) and non-ablated (B) animals on day 4 after ablation. Insets are higher magnifications of the animals in the boxed area. (C) Example of a double-headed polyp at day 12 after ablation. (D) Quantification of the formation of second heads. After ablation, all animals regenerated a second head (n = 20 for both ablated and non-ablated animals, 3 replicates). (E) Quantification of the regeneration of second heads after re-bisection at different time points. The percentage of double-headed animals gradually decreases as the interval between the first and second bisection increases.

Processing of Nifurpirinol by Nitroreductase is thought to eliminate the targeted cells by apoptosis [30]. In regeneration, apoptotic cells have different roles in different species, often being required for the regeneration process [1, 34, 35]. In *Nematostella*, blocking apoptosis has been shown to prevent head regeneration [36]. We therefore tested whether the unspecific induction of apoptosis by UV irradiation mimics the effect of ablating the *elav1^+^* neurons. We exposed juvenile polyps to increasing doses of UV, which caused 0 - 55% mortality 24 hours after exposure (Fig. S1). Following bisection of the irradiated animals, we did not observe any case in which a second head regenerated (Fig. S1). This suggests that induction of cell death by itself does not cause ectopic head formation.

### Ectopic head regeneration is preceded by upregulation of Wnt signaling genes

Wnt signaling controls head formation in *Nematostella* and other cnidarians [37] and overactivation of Wnt signaling has been shown to result in the formation of ectopic heads in *Nematostella* [23, 38, 39]. After bisection, Wnt signaling genes are upregulated specifically in head-regenerating tissue [27] and pharmacological activation of canonical Wnt signaling leads to ectopic head regeneration [23]. We therefore tested whether Wnt signaling genes are upregulated during the regeneration of ectopic heads in nervous system-ablated animals. As previously done for wildtype animals [27], we isolated the tips of regenerating head and foot fragments from ablated and non-ablated animals at different time points after bisection and analyzed the expression of selected genes by quantitative PCR. We focused on genes that displayed head regeneration-specific upregulation in a previous study [27]. To capture the changes in expression in the two conditions, we used the first timepoint of non-ablated head and foot fragments, respectively, for normalization of both ablated and non-ablated tissue. We denote this timepoint as 0-0.5 hours post amputation (hpa) because the procedure for tissue collection (bisection followed by removal of the regenerating tip) did not allow sampling immediately after bisection. *Wnt4* expression in the tip of foot fragments of non-ablated animals (i.e. fragments that regenerate a head) is upregulated between 6 and 24 hpa and returns to lower levels by 72 hpa (Fig. 4A). Expression after ablation followed the same pattern, though it remained upregulated at 72 hpa (Fig. 4A). In head fragments of non-ablated animals (i.e. regenerating a foot), *wnt4* expression decreased between 6 and 24 hpa, but overall, the changes were modest (Fig. 4B). In contrast, head fragments of ablated animals (i.e. regenerating a second head) showed elevated *wnt4* expression already at 0-0.5 hpa and further increased expression between 6 and 24 hpa (Fig. 4B), similar to the dynamics of head regeneration from foot fragments. We observed the same changes in the expression dynamics for *wntless*, a gene involved in secretion of Wnt molecules (Fig. S2).

**Figure 4.**
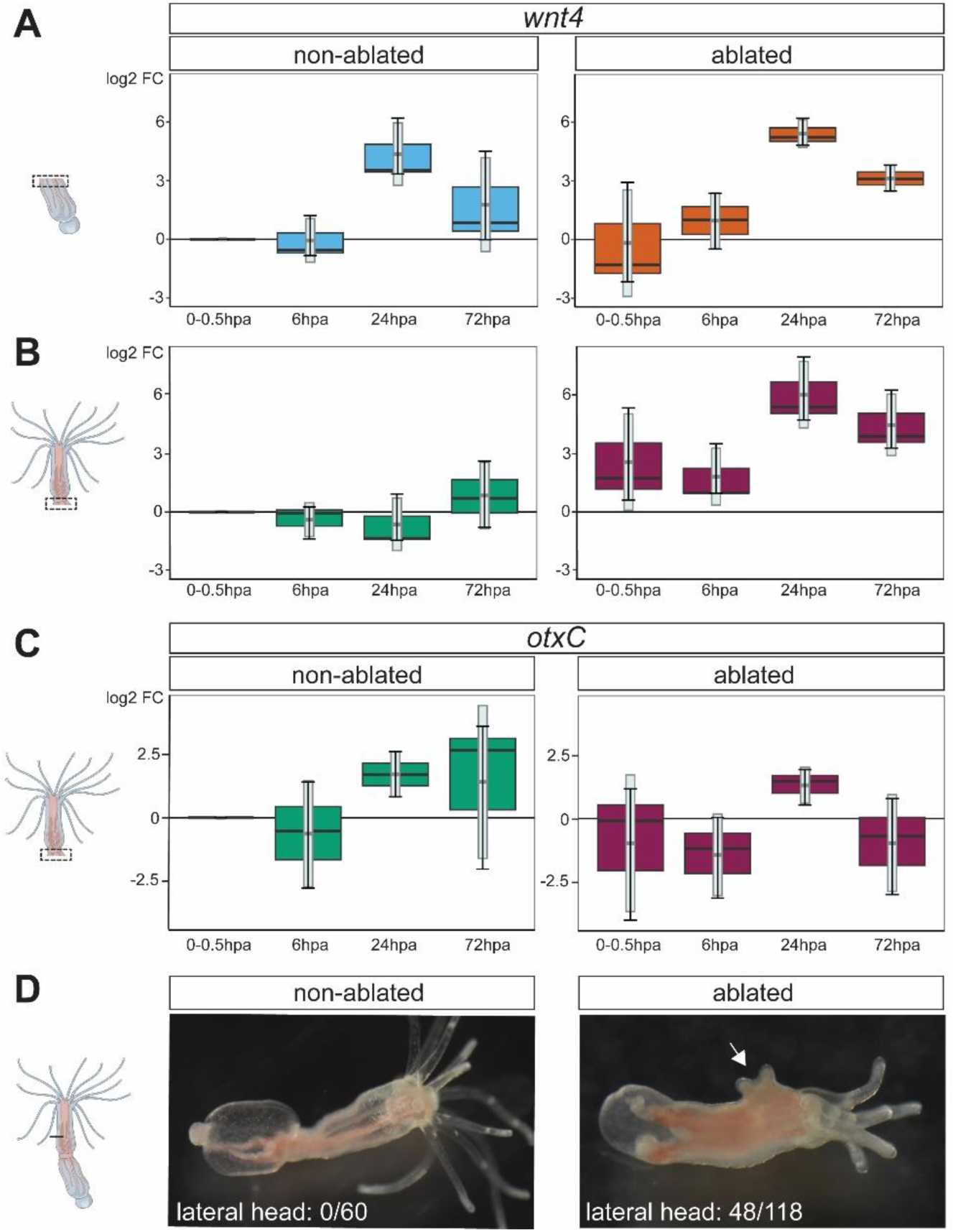
Early upregulation of Wnt signaling genes during ectopic head regeneration and formation of lateral heads after wounding. (A-C) Quantitative PCR for *wnt4* (A, B) and *otxC* (C) in non-ablated and ablated polyps. Cartoons on the left indicate the tissue used for cDNA synthesis. During ectopic head regeneration, *wnt4* expression is upregulated from the earliest sampling time point (B, right panel), whereas *otxC* is downreguated (C, right panel). Timepoint 0 – 0.5hpa (hours post amputation) of the non-ablated samples was used for normalization of the other timepoints of non-ablated, and all samples of ablated animals. Datapoints from the three replicates are visible as the median line in the box and as upper and lower whiskers. The light grey bars show the average +/- standard deviation. (D) Example of lateral head formation after wounding of ablated animals. Numbers represent the total of three replicates, with n = 20 per replicate for non-ablated, and n = 27-49 per replicate for ablated animals.

Expression levels of *otxC*, in contrast, increased during foot regeneration of non-ablated head fragments, both between 6 and 24 hpa, and between 24 and 72 hpa (Fig. 4C), as previously described [27]. In ablated head fragments, upregulation of *otxC* remains detectable between 6 and 24 hpa, but not at 72 hpa (Fig. 4C). Taken together, ectopic head regeneration after ablation of the *elav1*^+^ neurons and head regeneration in non-ablated animals employ an at least partially overlapping set of Wnt signaling genes.

### Formation of ectopic lateral heads after wounding of nervous system-ablated animals

Among the potential explanations for the regeneration of ectopic heads are (i) the local activation of a head regeneration-specific program at the aboral end of the head fragment, (ii) the inability to suppress a default head regeneration program caused by wounding [40, 41], or (iii) increased levels of head-like cues along the entire body column. In support of the latter scenario, we noted that the expression of *wnt4* and *wntless* was increased in the regenerating tips of ablated head fragments already at 0-0.5 hpa (Fig. 4B and S2). However, when we analyzed the expression of *wnt4* and *wntless* prior to bisection, we observed an only minimal increase in ablated compared to non-ablated animals. The expression of *otxC*, in contrast, was reduced (Fig. S3). As an alternative approach to test the propensity for generating ectopic heads, we inflicted small cuts in the upper body column. In non-ablated animals, these cuts always healed without signs of head formation. In contrast, following ablation of the *elav1*^+^ nervous system, ectopic heads formed at 38.3 ± 12.5% of the cuts (Fig 4D). Compatible with scenarios (ii) and (iii), this indicates that ablation of the *elav1*^+^ neurons leads to a generally increased potential for head formation, including in otherwise non-regenerative wounding.

## DISCUSSION

Here we show that ablation of a large part of the nervous system in *Nematostella* leads to the formation of a second head during regeneration from oral fragments. This phenotype was fully penetrant, indicating that the nervous system is required for suppressing a head regeneration program, but not for regeneration per se. The formation of ectopic head structures preceded the re-emergence of the first *elav1^+^* neurons and the ectopic upregulation of head regeneration-specific genes was apparent immediately after bisection of the ablated animals. We consider it, therefore, unlikely that cells of the nervous system directly regulate the identity of the regenerating tissue. Instead, we propose that the nervous system is required for establishing and/or maintaining graded positional information in the remaining tissue of the body wall and/or mesenteries. Such positional information might manifest in the level of Wnt signaling along the oral-aboral axis and the nervous system might be required to lower Wnt signaling in more aboral regions. The proposed signal from the nervous system could be chemical, i.e. via graded expression of secreted or cell surface molecules, or it could be based on differential or directional neural activity. The peristaltic constrictions of the body column, for example, move from oral towards aboral, likely reflecting directional neural activity. Silencing of the *elav1^+^* neurons without removing them might in the future allow addressing the potential role of neural activity in establishing or maintaining positional information along the body axis.

In the freshwater polyp *Hydra*, heating of the aboral end of head fragments leads to the regeneration of a second head in approximately 30% of the animals and this ectopic head regeneration can be blocked by a caspase inhibitor [42]. Apoptosis at the wound site occurs, however, both during head and foot regeneration and thus is unlikely to specifically instruct head regeneration [40, 43]. In *Nematostella*, apoptosis occurs at the wound site after bisection and blocking apoptosis prevents head regeneration [36]. The *elav1^+^* neurons are distributed rather uniformly along the body column [29], and their ablation thus does not result in apoptosis at a specific site along the oral-aboral axis. Furthermore, global induction of cell death by UV irradiation did not lead to the formation of second heads after bisection. It is therefore unlikely that apoptosis as such promotes secondary head formation, but we cannot rule out that a neuron-specific aspect of apoptosis contributes to the proposed increase in positional information throughout the polyps.

There are numerous examples for an involvement of neurons in various aspects of regeneration and in many different species [44–46]. For whole-body regeneration, a possible role of the nervous system has been addressed in some other highly regenerative species. In *Hydra*, the removal of interstitial stem cells eventually leads to the absence of neurons, nematocytes (stinging cells) and most gland cells. Such neuron-free animals regenerate with the correct polarity and they do not display signs of altered positional information in transplantation assays [16, 17]. In planarians, a contribution of the nervous system in polarity control has been suggested by experiments in which simultaneously disrupting the ventral nerve cord and blocking gap junctions resulted in ectopic anterior regeneration at posterior wounds [47]. Furthermore, silencing of the nervous system-specific gene *roboA* caused the regeneration of ectopic pharynx tissue with inverted polarity [48]. But, to our knowledge, ablation experiments specifically addressing the role of neurons have not been reported for planarians. Since positional information in planarians is present in subepidermal muscle cells [14], a scenario in which the nervous system affects this positional information - akin to the role proposed here for *Nematostella* - appears plausible. In some annelid species, the presence of the ventral nerve cord at the wound site is required for regeneration [49–53] and diverting the nerve cord to a lateral wound is sufficient to induce ectopic head formation in other species [54]. However, systematic regeneration of an incorrect structure (head or tail) as a consequence of nervous system removal has – to the best of our knowledge - not been described. These examples illustrate that the data are currently too fragmented and the experimental paradigms too disparate to conclude whether an involvement of the nervous system in determining polarity during whole-body regeneration is shared among different taxa.

Taken together, we identified a requirement for the nervous system in controlling tissue polarity during whole body regeneration in *Nematostella* by suppressing a default head regeneration program. These findings open the door for studying the possible evolutionary conservation of this function by directly comparable experimental approaches in other highly regenerative and genetically tractable species [55–59].

## MATERIAL AND METHODS

### Animal culture

Adult polyps (derived from CH2 x CH6 [60]) were maintained in diluted seawater (*Nematostella* medium, 12-14ppt) at a temperature of 18-19°C and fed with freshly hatched *Artemia* nauplii. Spawning was induced by a temperature shift and exposure to light as described in [61]. The trangenic line *elav1::mOrange* is described in [29], *MyHC::mCherry* in [32], and *foxQ2d::mOrange* in [31].

### Generation of the transgenic line

The *elav1::nitroreductase-cerulean* transgene is based on *elav1::mOrange* [29]. The ORF for mOrange was replaced by *cerulean* lacking the start codon. The coding sequence of *nitroreductase* was amplified from *E.coli* and inserted in front of *cerulean* using AscI sites. Meganuclease-mediated transgenesis was carried out as described previously [62, 63], polyps hemizygous for *elav1::mOrange* or *elav1::ntr-cerulean* were crossed to obtain double transgenic animals.

### Nifurpirinol treatment and bisection

After fertilization, animals were kept at 21°C in the dark, and they were fed five times per week from 10 days post fertilization until an age of six-seven weeks. Nifurpirinol (Merck 32439) was applied for 72 hours at a final concentration of 10µM in *Nematostella* medium + 0.5% DMSO, the solution was exchanged once per day. During the exposure, animals were kept at 25°C. At the end of the treatment (at D0), the animals were washed 3x with *Nematostella* medium, transferred to fresh dishes and returned to 21°C. For determining the time course of nervous system regeneration (shown in Figure 1), the animals were fed daily starting from day 3 after washout of Nfp. For bisection and horizontal cuts, polyps were relaxed with MgCl_2_, the bisections/cuts were done below the pharynx and after bisection the oral and aboral ends were separated. The animals were not fed during the bisection experiments shown in Figure 3.

### UV treatment

Animals in *Nematostella* medium were placed in open petri dishes and exposed to UV light (254nm) in a BLX-E254 crosslinker (Vilbert). Bisections below the pharynx were done on the following day.

### RNA extraction, reverse transcription and real time qPCR

The regenerating tips of head and foot fragments, respectively, were collected at different time points after bisection and placed in 1 ml TriReagent (Sigma), incubated at room temperature for 5 min and subsequently frozen at -80°C. After thawing, the samples were vortexed, chloroform (200 µl) was added and the samples were vigorously mixed. After incubation at room temperature for 1 min, samples were centrifuged at 12000 g and 4°C for 15 min. The upper phase was transferred to a new tube and 500 µl of Phenol-chloroform-isoamyl alcohol mixture (Sigma) was added. After mixing and incubating at room temperature for 2 min samples were centrifuged at 12000 x g at 4°C for 10 min. To the upper phase 500 µl of chloroform was added, samples were mixed and incubated at room temperature for 1 min. Centrifugation was performed at 12000 x g for 10 min at 4°C. To the upper phase RNA grade glycogen (Thermo Fisher Scientific R0551) and 500 µl isopropanol were added. After mixing and incubation at room temperature for 20 min samples were centrifuged at 13000 x g for 20 min at 4°C. The pellet was resuspended in ethanol (70% ice cold) and centrifuged at 8000 x g for 5 min at 4°C. The pellet was dissolved in RNAse-free water. The quality and concentration of the RNA was checked by Epoch Spectrophotometer (Agilent). The High-Capacity cDNA Reverse Transcription Kit (Thermo Fisher Scientific 4368814) with RNase Inhibitor was used to generate cDNA according to the manufacturers protocol.

Real-time qPCR was performed from three biological replicates using PowerUpTM SYBRTM Green Master Mix (Thermo Fisher Scientific A25918). The reaction mix contained 10 µM of each primer and 1 ng of cDNA. 39 cycles were performed with 60 °C as the annealing step. The primer efficiency for each primer set was measured using a standard curve based on five serial cDNA concentrations, expression was normalized to *Hsp70* and calculated using the C_T_(n^-ΔΔCt^) method. Primers are as in [27], they are listed in Table S1.

### Electron microscopy

Polyps were relaxed for 30 min in 7% MgCl_2_ and *Nematostella* medium mixed 1:1 and then fixed in 2.5% glutaraldehyde in 0.1M PBS. They were postfixed in 1% Osmium tetroxide in the same buffer, dehydrated in a graded acetone series and embedded in Epon/Araldite. All steps were performed in a microwave (Pelco BioWave®Pro +, Ted Pella, USA). Sections of 50 nm were cut with an ultra 35° diamond knife (Diatome, Switzerland) on a UC7 ultramicrotome (Leica) and collected on Beryllium-Copper slot grids (Synaptek, USA) coated with 1% polyetherimide (in chloroform) and contrasted with 2% uranyl acetate (in water) and lead citrate. Sections were imaged with STEM-in-SEM [64] at a resolution of 25 and 11 nm/pixel with a Zeiss Supra 55VP and with Zeiss SmartSEM®.

### Confocal microscopy

Animals were relaxed in MgCl_2_ and DNA staining was performed with Hoechst 33342 (Thermo Fisher Scientific) in *Nematostella* medium (final concentration 100µg/ml). Live animals were placed on glass slides while in the MgCl_2_ solution. Imaging was performed with OLYMPUS FV3000 or Leica TCS SP5 confocal laser scanning microscopes. Imaging files were managed in FIJI [65], figures were assembled in Adobe Illustrator.

### Analysis of peristaltic waves and contraction

For observing peristaltic waves, animals were recorded for five minutes. The poking assay was performed as previously described [66]. Briefly, once the individual was relaxed a glass pipette was used to lightly touch the oral opening of the animal while observing under a dissection scope. Animals were given up to three chances to respond to the touch by retracting their tentacles and/or body column. To be considered responsive, the animal had to react in at least one out of the three pokes performed. The same animals were recorded pre- and post-ablation to determine how the loss of *elav1^+^* neurons affected the ability to respond to a touch stimulus.

## ACKNOWLEDGEMENTS

We thank Eilen Myrvold, Brandon Reid Mellin and Lavina Jubek for excellent care of the *Nematostella* facility at the Michael Sars Centre and the ELMILAB facility at the University of Bergen for access to electron microscopy. We thank Eric Roettinger, Aldine Amiel, Chiara Sinigaglia, Marta Iglesias and members of the Rentzsch group for very helpful discussions, and Kathrin Garschall for advice on UV experiments. Work in FR’s lab was supported by grant 251185/F20 from the Research Council of Norway and the University of Bergen.

