## Supplemental Information for "Ectopic head regeneration after nervous system ablation in a sea anemone"

**for**

| | Replicate 1 | | | | Replicate 2 | | | |
|---|---|---|---|---|---|---|---|---|
| UV [J/cm$^2$] | n | survival 24h | survival 8d | ectopic head | n | survival 24h | survival 8d | ectopic head |
| 0.05 | 40 | 40 | 37 | 0 | 40 | 35 | 35 | 0 |
| 0.1 | 40 | 36 | 34 | 0 | 40 | 31 | 27 | 0 |
| 0.2 | 40 | 27 | 25 | 0 | 40 | 29 | 27 | 0 |
| 0.3 | 40 | 35 | 26 | 0 | 40 | 22 | 16 | 0 |

**Figure S1 UV irradiation does not cause ectopic head regeneration.**

Following exposure to the indicated doses of UV, animals were allowed to recover for 24h and then bisected below the pharynx. All surviving head fragments closed the wound, but none regenerated a second head.

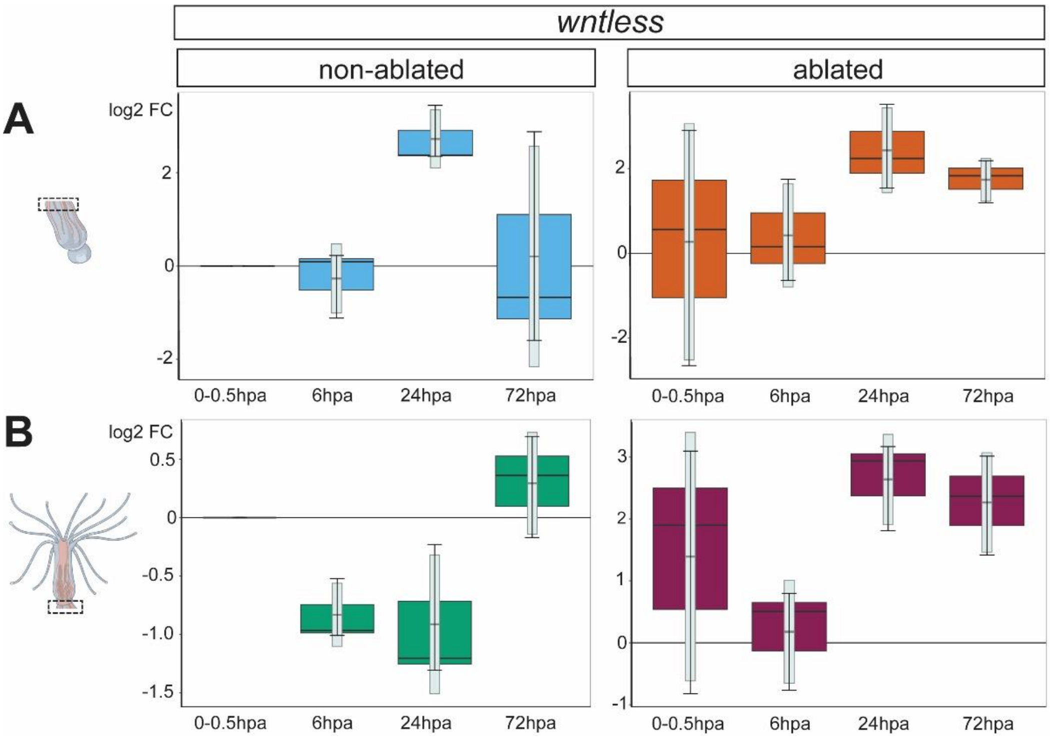


**Figure S2 Expression of *wntless* is upregulated during ectopic head regeneration.**

Quantitative RT-PCR for *wntless* in regenerating tips of (A) foot and (B) head fragments in either ablated or non-ablated animals. Similar to *wnt4*, *wntless* transcription is upregulated during the regeneration of ectopic heads. Datapoints from the three replicates are visible as the median line in the box and as upper and lower whiskers. The light grey bars show the average +/- standard deviation.

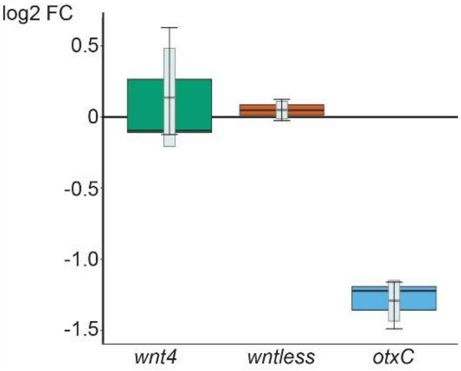


**Figure S3 Gene expression levels in ablated vs non-ablated polyps.**

Quantitative RT-PCR of *wnt4*, *wntless* and *otxC* in ablated vs non-ablated animals after washout of Nfp (day 0). Values for non-ablated animals are defined as 0. Only *otxC* is differentially expressed. Datapoints from the three replicates are visible as the median line in the box and as upper and lower whiskers. The light grey bars show the average +/- standard deviation.

| Oligo Name | Sequence (5'-3') | Source |
|---|---|---|
| Nv-wnt4_fwd<br>Nv-wnt4_rev | TAACAGAGGCGGAAGACTGG<br>ATGTAGTCGCCGACGATTCT | [27] |
| wntless_fwd<br>wntless_rev | CTCACCCTGGGCTAGAGATG<br>AAGTCGTGCTTCAAACGTCC | [27] |
| otxC Fw<br>otxC Rev | GCCCACTGCGACTTTTCTAT<br>GCTCGTTCTTGCTCGGTATC | [27] |
| Nv-Hsp70_fwd<br>Nv-Hsp70_rev | TCGATGATCCTGGGGTAAAG<br>CCTGCCTCGTTCACTACCTC | [27] |

**Table S1 Primers for quantitative RT-PCR**

**Video captions**

**Video S1: Feeding after ablation of the *elav1*$^{+}$ neurons.**

The video recording is shown at real-time speed.

**Video S2: Peristaltic waves in a non-ablated polyp.**

The video recording is shown at 6x real-time speed.

**Video S3: Contraction in response to poking in a non-ablated polyp.**

The video recording is shown at 6x real-time speed.

**Video S4: Contraction in response to poking after ablation of the *elav1*$^{+}$ neurons.**

The video recording is shown at 6x real-time speed.
